# Accuracy of different modalities of reaction time testing: Implications for online cognitive assessment tools

**DOI:** 10.1101/726364

**Authors:** Jameson Holden, Eric Francisco, Rachel Lensch, Anna Tommerdahl, Bryan Kirsch, Laila Zai, Robert Dennis, Mark Tommerdahl

## Abstract

Reaction time testing is widely used in computerized cognitive assessments, and clinical studies have repeatedly shown it to be a sensitive indicator of cognitive function. Typically, the reaction time test is administered by presenting a subject with a visual stimulus on a computer monitor and prompting the individual to respond (via keypad or computer mouse) as quickly as possible. The individual’s reaction time is calculated as the interval between presentation of the stimulus and the time recorded from the mechanical response. However, there are many inherent latencies and variabilities that may be introduced to the measure by both hardware (computer monitor and mouse) and software (operating system). Because of these delays, we hypothesized that a comparison of hardware protocols (excluding human response) would demonstrate significant differences in the resulting reaction time measures. To simulate the delays of various components of the common systems used to obtain reaction time, we conducted a simple experiment in which either a visual or tactile stimulus evoked a movement from a mechanical transducer to respond to a computer peripheral or a dedicated response device. In the first condition, a simulated visual reaction time test was conducted by flashing a visual stimulus on a computer monitor. The stimulus was detected by a dedicated light sensor, and a linear actuator delivered the mechanical response via computer mouse. The second test condition employed a mobile device as the medium for the visual stimulus, and the mechanical response was delivered to the mobile device’s touchscreen. The third and fourth test conditions simulated tactile reaction time tests in which the stimulus was generated by a dedicated hardware device. The third condition simulated a tactile stimulus, which was detected by a mechanical switch, and again a hardware device delivered the response via computer mouse. The fourth condition also simulated a tactile stimulus, but the response was delivered by a dedicated hardware device designed to store the interval between stimulus delivery and stimulus response. There were significant differences in the range of responses recorded from the four different conditions with the reaction time collected from a visual stimulus on a mobile device being the worst and the device with dedicated hardware designed for the task being the best. The results suggest that some of the commonly used visual tasks on consumer grade computers could be introducing significant errors for reaction time testing and that dedicated hardware designed for the reaction time task is needed to minimize testing errors.

## Introduction

A study by Woodley et al. (2013) postulated that as a human race, we are getting “dumber”. The basic premise of the study was that reaction times are getting slower, and that this contradicts a number of other studies that had demonstrated that, based on performance on IQ tests, we are actually getting a bit smarter. The purpose of this report is not to weigh in on whether or not humans are getting dumber as a species, but rather to focus on the accuracy of reaction time testing and how it has changed historically.

There have been many reaction time studies over the past 150 years. The graph in Figure 1 is a summary of the data points obtained from healthy subjects across a collection of those publications. The data demonstrate not only an upward drift of reaction time, but a larger range of reaction times, with the progressive degradation of reaction time appearing to begin in the 1970s and 1980s. Thus, the question that the authors think should be asked is not whether we are getting worse at reaction time testing, but could there be inconsistencies introduced by the reaction time testing itself?

Note that while early reaction time studies (between 1850 and 1950) demonstrated human performance in the 150-200 msec range, the range reported post-2000 extends from 150 to 400 msec. This enormous shift suggests either a very different population that is being tested or a very different strategy for measuring reaction time. Since the dumbing down of the human species is really not something that the authors believe can be accurately measured (other than observations of how much time some generations spend utilizing social media), we targeted addressing the actual methods that were being used. Casual observation of the data plotted in Figure 1 suggests that the methods underlying the represented scientific literature are becoming much more variable and consequently, less accurate.

**Figure 1.**
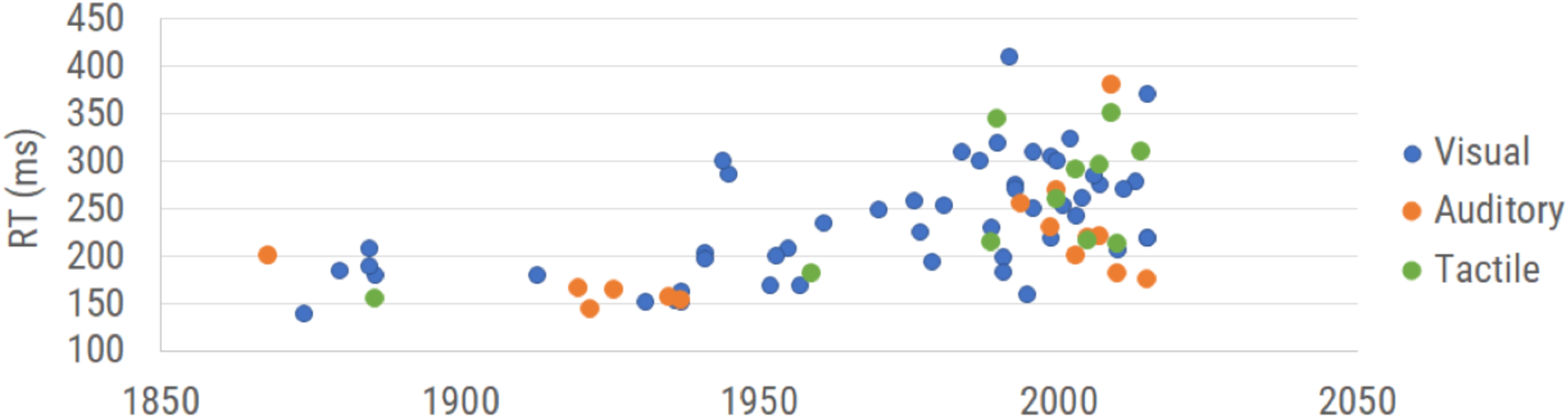
Results of reaction time testing reported in the literature. Reaction time values are categorized by modality of measurement (visual reaction time, auditory reaction time, and tactile reaction time) and plotted in the corresponding year collected.

Reports in the literature describing and/or utilizing reaction time tests date back to the 19th century. In 1849, Hermann von Helmholtz used electric stimulation to investigate nerve conduction velocity, first examining the conduction velocity in the legs of frogs before modifying his methods to accommodate human subjects. Helmholtz stimulated the skin at two separate locations and measured the time required for the human subject to respond via hand signal to the stimulation at each location. By measuring the distance between the two points of stimulation and the difference in their associated reaction time, he deduced a fairly accurate estimate of nerve conduction velocity. Donders expanded on the concept to include central as well as peripheral nervous system processing in which he stimulated either skin, eye or ear and had the subjects respond with their hands (Donders 1868). Time recorded was based on a chronograph, and although the early methods may seem crude or cumbersome by contemporary methods, the results from these early experiments inspired numerous investigations utilizing reaction time over the subsequent 150 plus years. The methods used by Merkel, visual stimulation and tactile response (Merkel, 1885), are still commonly used due to the simplicity of implementation, though it is noteworthy that the individual data obtained by Merkel ranged between 152 and 201 msec, which is not the case for most contemporary studies.

Many of the studies spanning the 150-year time frame investigated changes in reaction time that resulted from a number of neurological disorders or insults. For example, reaction times have been demonstrated to be altered by changes introduced by neurological insults such as TBI/mTBI (Ruesch, 1944; Van Zomeren and Deelman, 1978; Macflynn et al., 1984; Stuss et al., 1989; Ponsford and Kinsella, 1992; Collins and Long, 1996; Zahn and Mirsky, 1999; Warden et al., 2001; Collins et al., 2003; Sarno et al., 2003; Willison and Tombaugh, 2006; Niogi et al., 2008; Gould et al., 2013; Eckner et al., 2015), PTSD (Ruesch, 1944), pharmaceuticals (Edwards and Cohen, 1961; Ancelin et al., 2006), aging (Benton, 1977; Sherwood and Selder, 1979; Fozard et al., 1994; Lajoie and Gallagher, 2004; Der and Deary, 2006), dementia (Ancelin et al., 2006), Parkinson’s (Evarts et al., 1981; Goodrich et al., 1989), schizophrenia (Schwartz et al., 1989), ADHD (Meere et al., 1992; reviewed in Tamm et al., 2012; Puts et al., 2017), sleep deprivation (Lorenzo et al., 1995), caffeine (Cheney, 1934), alcohol (Hernandez et al., 2006), autism spectrum disorders (Puts et al., 2014; Ferraro et al., 2014), and diabetes (Patil and Phatale, 2015). The widespread utilization of the reaction time test across many decades of research and its utility in many different clinical and clinical research venues led us to ponder how the accuracy of this measure might have changed historically. Contemporary users of the reaction time metric might assume that using modern and faster computer technology automatically leads to more accurate reaction time measures. The fallacy of this assumption is that the modern computer technology commonly deployed for reaction time testing is not designed to be laboratory equipment, which causes the accurate timing of external events to suffer. Laboratory equipment of the mid-19^th^ century was specifically designed for laboratory use and was probably as accurate at performing reaction time testing as many of today’s computer based reaction time tests, if not more so.

The question we sought to address was whether or not modern computing methods introduce problems to the reaction time measure. There are inherent delays predicted to be introduced by both software and hardware. Many contemporary reaction time tests are administered through a computer program that calculates the time elapsed between stimulus delivery and the subject response (typically the click of a mouse). The majority of these tests use either a visual (e.g. screen flash) or auditory (loud beep) stimulus, as these stimuli can be delivered using commodity-grade human-computer interfaces such as computer monitors, mice, keyboards, and touch screens. Reaction time tests that employ a different mode of stimulation (such as a tactile stimulus) require additional hardware, which may be connected to the test computer by a physical or wireless connection. The reaction time test is contingent on the CPU timing accuracy of the testing computer, which can vary based on which programs are running in the background and the inherent processing speed of the chip. Also of great concern is the operating system (OS) timing cycle and task priority structure, which typically manages many tasks, including system overhead unrelated to the reaction time test in progress. While this division of attention is seldom apparent to the user, it typically introduces delays of ∼15 ms, or more when the computational demand is high, such as in the presence of malware or background tasks, or as a result of widely employed networking prioritization such as “audio prioritization”. In this case, the delays in other functions can even become clearly apparent to the user, being on the order of hundreds of milliseconds or more. Even at its minimum this CPU latency is typically between 2-20 ms, which will significantly and ambiguously alter reaction time test results. In addition to OS latency, different device driver firmware can introduce latencies differing by tens of milliseconds or more between drivers, even with identical hardware (Plant and Turner, 2009). These computer hardware and software variations can introduce variable timing delays of up to 100 msec (Neath et al, 2011).

Reaction time tests suffer from latencies that are introduced at points aside from core processor timing in their protocols as well. For example, these can be introduced by the commodity human interface peripherals. USB and wireless mice and keyboards introduce latency in their communication protocols at several points, including pre-transmission buffering, transmission, and post-transmission buffering before transfer to the CPU for processing. Most computer screens have a refresh rate of 60 Hz, and a screen flash can occur up to 17 ms before or after the ‘stimulus delivery’ time that is recorded by the CPU. Touch screens on both mobile and desktop devices have a built-in latency related to the sensing mechanism, usually by capacitance. For smooth operation, touch sensing requires a certain amount of signal processing both in hardware and software or firmware, because touch signals typically involve a certain amount of “integration” at or near the sensor. This is necessary both for noise mitigation, to eliminate spurious signals, as well as to detect the proximity of a finger or stylus by capacitive coupling. For example, capacitive sensing can be done in several different ways, but these all invariably involve the rate of charge or discharge of a capacitor which is modified by the proximity of a finger or stylus that changes the value of the capacitor being charged, and thus the RC time constant, by a few percent. Styli may be standardized for a specific device, but fingers are not, and thus it is necessary to set thresholds and make decisions in firmware whether or not a touch event has occurred, or not, for a wide range of contact conditions. All of this integration and processing takes time, even when done by distributed processing. The same arguments can be modified and applied to older force sensing screen interfaces which had the added computational burden of calculating a force centroid to determine where force was being exerted on the plane of the display, and similarly for any other touch sensing strategy such as resistive or others. Layered on top of this is the firmware task of interpreting what type of touch is being detected, whether there are one or multiple touch points. Once the signal has been cleaned, filtered, and interpreted by the peripheral touch sensing device, it can then be placed in the communication buffer, where it waits its turn for CPU priority. Because of all the variables involved, and different strategies employed by touch sensing peripheral hardware, it is not possible to calculate the hardware latency. As a result, current “lag” from touch screen-to-display varies from 50 ms to 200 ms (Ng et al. 2012).

The objective of this study was to determine if there are significant and measurable differences introduced to reaction time measures that are collected with different types or categories of hardware currently used in commercially available reaction time tests. More specifically, how would a visual reaction time test performed on a computer laptop or mobile device with consumer grade hardware (most common method) compare with a tactile reaction time test delivered with laboratory grade hardware (less commonly performed). If equipment and/or operating systems do in fact introduce errors into the reaction time test, then it would be hypothesized that different methods requiring significantly different hardware or software would generate very different reaction times and reaction time variability for individuals taking the reaction time test. In order to directly investigate the differences introduced by various testing strategies, the human element was removed from this study and automated/dedicated hardware was used to perform the reaction time tasks. Four modes of reaction time testing were evaluated robotically to compare the potential contributions of different user interfaces to the reaction time test.

## Methods

Experiments were conducted with four different conditions in order to observe results obtained with variable stimulus and response protocols for a reaction time test with a non-human interface. The stimuli delivered were visual/optical (simulation of a visual stimulus) with a mobile device, visual/optical with a computer monitor, and tactile/mechanical (simulation of a mechanical stimulus) with a dedicated hardware device (mechanical stimulus delivered with the Brain Gauge; Cortical Metrics, LLC). The response methods utilized were tactile/mechanical via touchscreen (simulation of finger press on touch screen), tactile/mechanical via computer mouse (simulation of finger response via computer mouse), and tactile/ mechanical via dedicated hardware device (simulation of finger response via Brain Gauge).

### Apparatus and device setup

Four simulations were performed and different configurations were used to deliver those simulations. In each case, simulation of an individual taking a reaction time test was performed by detection of a visual or mechanical stimulus via electronic switch to simulate stimulus detection and delivery of a mechanical stimulus to simulate a finger depressing a response device. The configurations were assembled with a standard breadboard. Each of the following tasks ran for N=100 trials and each simulation was conducted with the same CPU (MacBook Pro 2017). Trials were performed for either a visual or mechanical stimulus simulation. Detection simulation was performed by mechanical response to either a computer mouse, a touchscreen or a dedicated device (Brain Gauge; Cortical Metrics, LLC).

### Simulation #1: Visual stimulus & response via computer mouse

An analog light sensor (Adafruit ALS PT19) was placed 2 mm in front of an LCD monitor (60 Hz, 1080p, Dell) and was programmed to flash from black to white at random intervals (4-6 seconds). The light sensor was configured with a triggering threshold of 800 mV. In order to simulate the response interval of a reaction time test, a dedicated mechanical apparatus was designed that was triggered to respond automatically at a fixed interval (100 ms) after the flash was detected by the light sensor. The mechanical apparatus (Figure 2) was assembled on a standard breadboard and was comprised of a linear voice coil actuator (VCA), a 555 Timer configured for monostable output (110 +/-1 ms), an N-channel field-effect transistor (FET), and a 5V power supply.

**Figure 2.**
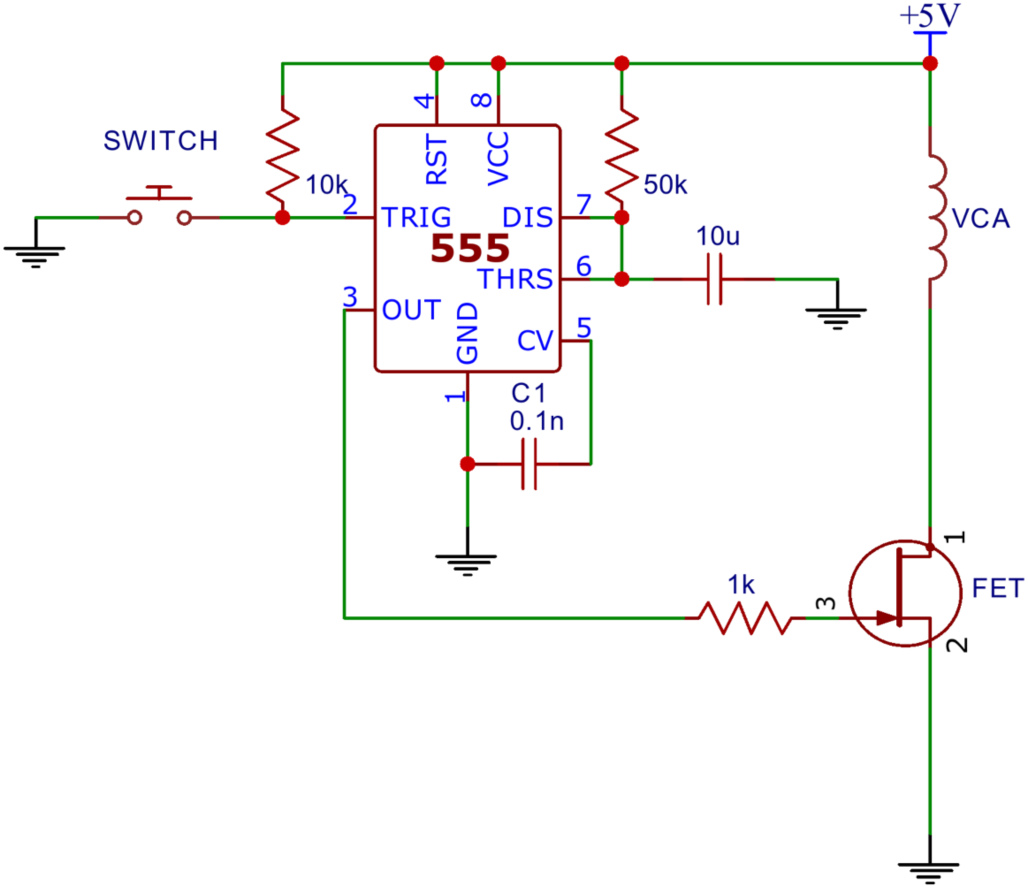
Dedicated mechanical apparatus.

The VCA of the mechanical apparatus was positioned directly above the left button of the USB computer mouse in order to simulate a button press by a human (Protocol 1 of Figure 3). After the programmed delay the VCA would receive a 10 ms pulse from the 555 timer which caused the VCA to depress the mouse button by 800 microns. The mouse button reached its triggering point at the middle point of the pulse, theoretically adding 5 ms to the 100 ms programmed delay. Thus, the expected true reaction time for the system was 105 ms.

**Figure 3.**
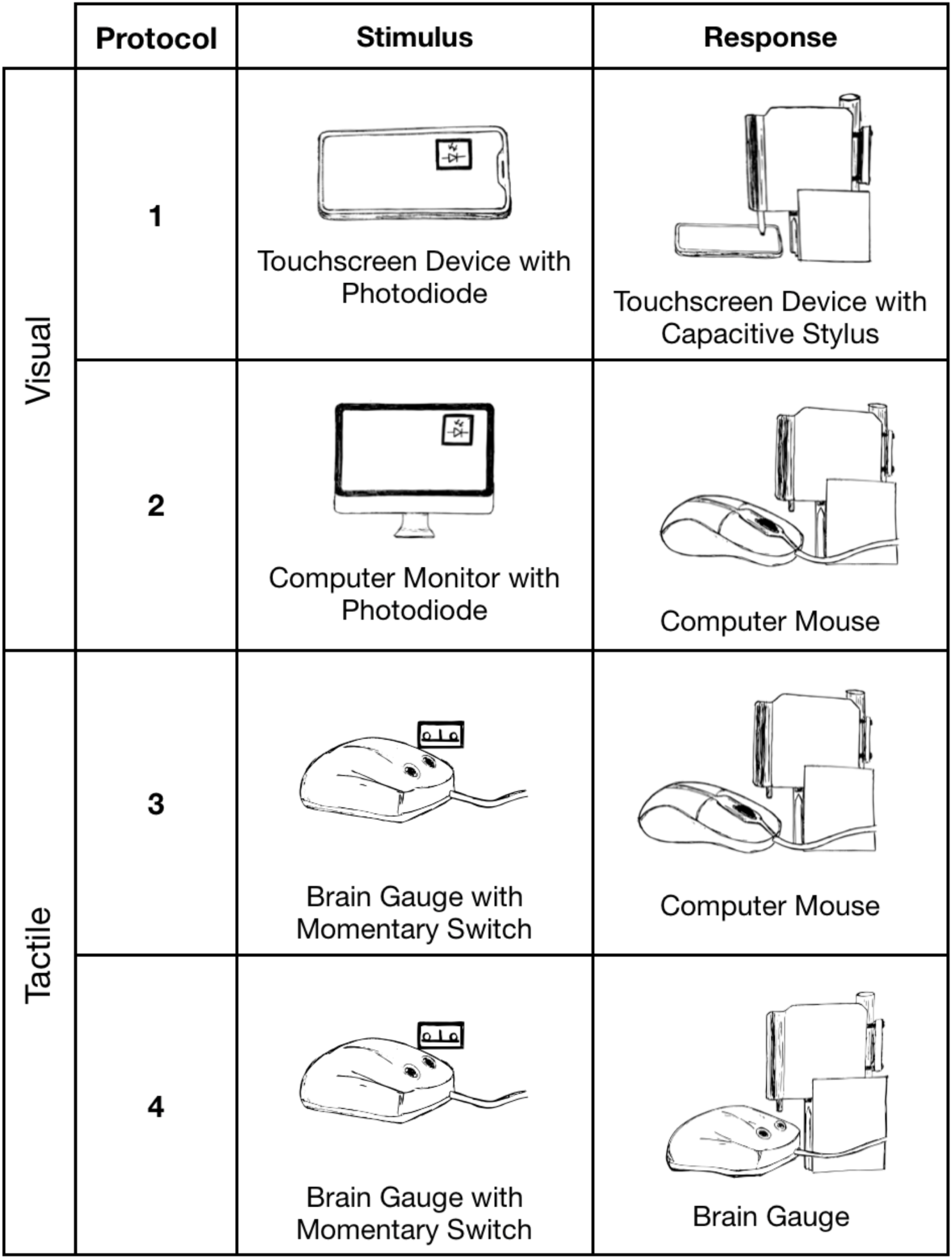
Reaction time simulations. Protocols 1 and 2 delivered mechanical stimuli in response to a visual stimulus. Protocols 3 and 4 delivered mechanical stimuli in response to a tactile stimulus.

### Simulation #2: Visual stimulus & response via mobile device

The analog light sensor was mounted 2 mm above a mobile touchscreen (Nextbit Robin, Android 7.1.1 “Nougat”), which was programmed to flash from black to white at random intervals (4-6 seconds). The light sensor was configured with a triggering threshold of 800 mV. A capacitive sensor was mounted to the VCA probe tip of the apparatus and placed 2.6 mm above the touchscreen to simulate a subject’s response. The mechanical response from the VCA was adjusted to deliver a tap with an amplitude of 2.6 mm over a duration of 10 ms to the touchscreen. The mechanical response was simulated as in Simulation #1 and was used to simulate a finger press 100 ms after the flash triggered the light sensor. Expected true reaction time for the system was 105 ms.

### Simulation #3: Tactile stimulus & response via computer mouse

A mechanical switch (Cherry MX Red) was mounted above the probe tip of a tactile stimulator (Brain Gauge; Cortical Metrics, LLC) in order to detect a mechanical stimulus of a simulated reaction time test. A 1.5 mm stimulus was used to depress the switch above the actuation point and trigger the mechanical response simulator circuitry. The VCA probe tip of the mechanical apparatus was positioned above a computer mouse to simulate a subject’s responding digit in a resting state. The stimulus pattern, programmed delay and switch response were identical to the conditions in Simulation #1. The mechanical apparatus from Simulation #1 was modified to simulate a controlled human reaction time of 100 ms after the mechanical switch was triggered. Expected true reaction time for the system was 105 ms.

### Simulation #4: Tactile stimulus & response with dedicated hardware device

The mechanical detection system was arranged as in Simulation #3. The VCA probe tip was mounted above the response tip of the dedicated hardware reaction time device, depressing the device’s tip by 1.5 mm. The VCA response was identical to the task of Simulation #3. The mechanical apparatus from Simulation #3 was used to simulate a controlled human reaction time of 100 ms after the mechanical switch was triggered. Expected true reaction time for the system was 105 ms.

## Results

Four simulated reaction time tests were performed. In each case a stimulus (visual or tactile) was delivered and detected electronically, and a response was made mechanically either via touchscreen, USB mouse or with a dedicated testing device (Brain Gauge; Cortical Metrics, LLC). The expected true reaction time for all four experimental groups was 105 ms; Figure 4 displays the results.

**Figure 4.**
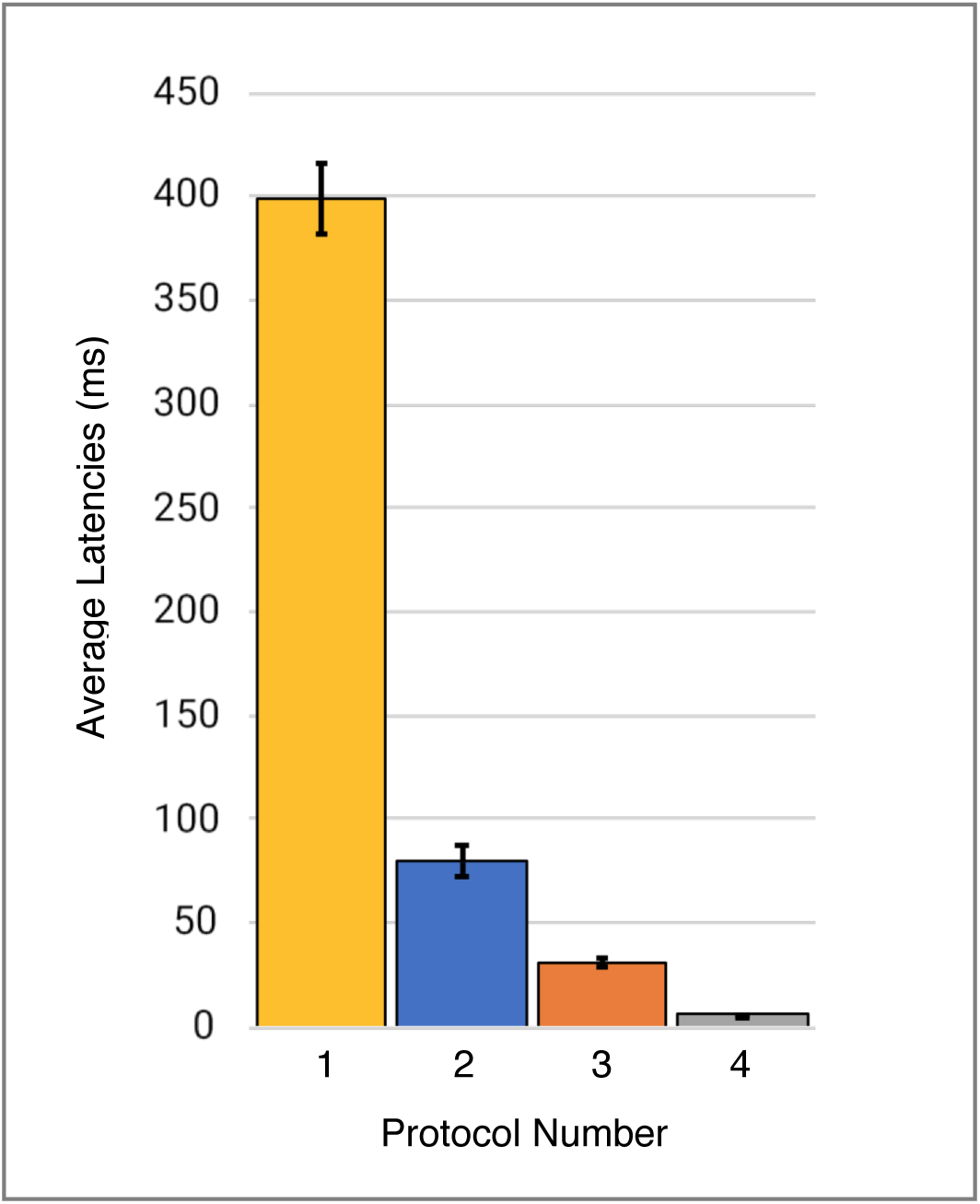
Comparison of the accuracy of four different methods of reaction time testing. Numbers on horizontal axis correspond to protocols described in Figure 1. (1) Visual stimulus with response on mobile device (yellow) has the highest value, was the least accurate and has the largest variability. (2) Visual stimulus with response on a computer mouse (blue) was the second highest. (3) Tactile stimulus with response on a computer mouse (orange) was significantly better than either visual modality although not as accurate as (4) tactile stimulus with dedicated tactile hardware response device (gray).

Reaction time to a simulated visual stimulus in which a touchscreen was used as the response device generated the highest latency of 400 msec. When the same visual stimulus simulation was coupled with a response from a USB Mouse, reaction time latency was significantly improved to 80 msec. Reaction time to a tactile stimulus simulation utilizing the same USB mouse for a response device demonstrated a latency of 30 msec. Reaction time to the tactile stimulus simulation with response on the dedicated tactile device had the smallest latency error. Average latencies are plotted in Figure 4.

Perhaps more significantly than the latency on each task was the variability. In Table 2, note the variability and range of latencies. This variability is prominently noticeable in Figure 5 which displays the data point-by-point.

**Figure 5.**
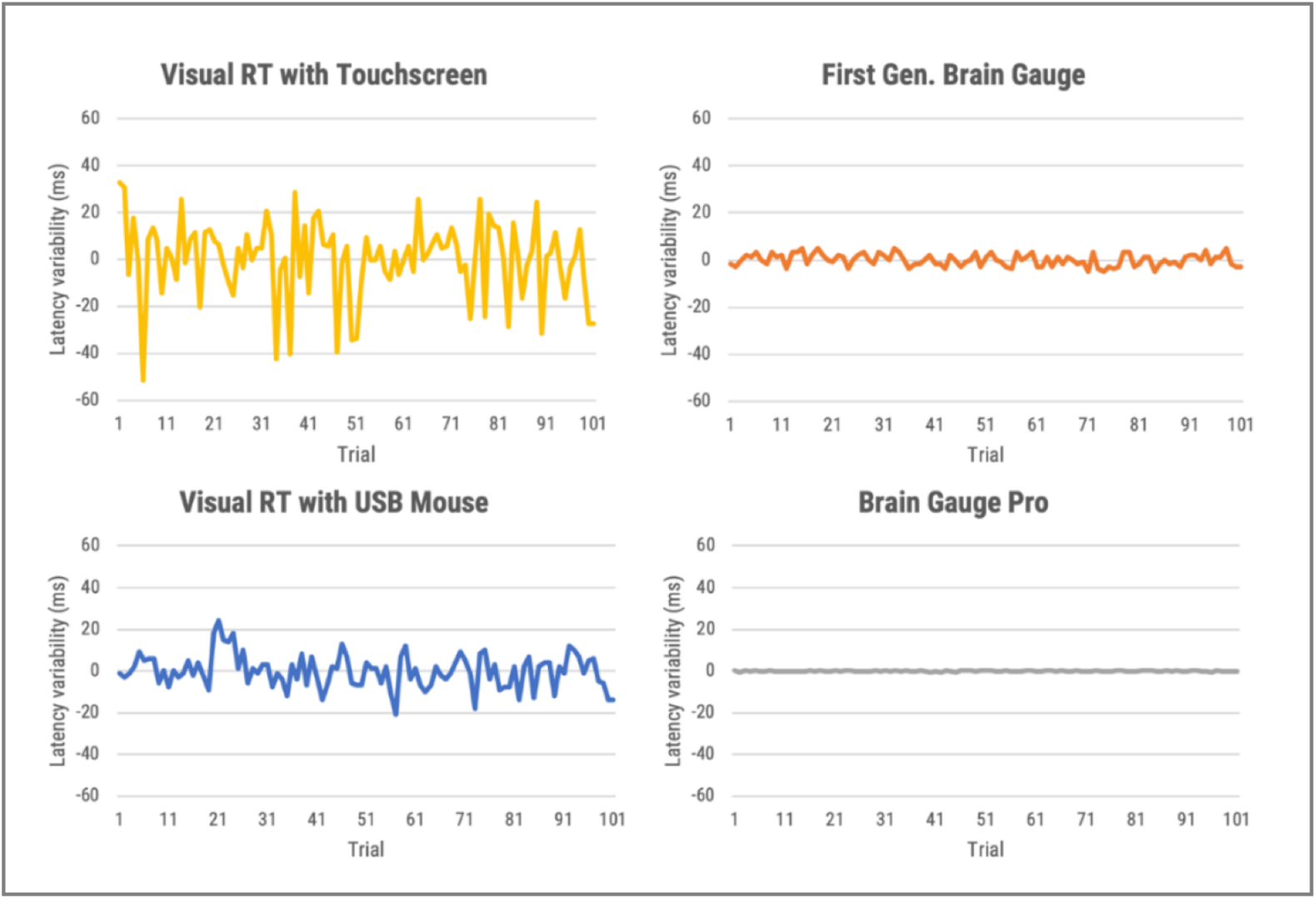
Direct comparison of data from the four reaction time testing methods with averaged offset subtracted. Raw data is plotted with offset of median latency subtracted, a technique used by many reaction time assessments to adjust for systematic latencies. All data is plotted on the same scale. The visual task simulations had significantly greater variabilities than the tactile simulations.

## Discussion

In this study, we demonstrated that reaction time testing, simulated robotically, shows profound differences in performance when stimuli are delivered either by visual or tactile modalities, and that there are significant differences in performance when the responses to those stimuli are delivered mechanically via either touchscreen, USB mouse, or a dedicated device that was designed for the task. In other words, a comparison was made in reaction time performance between consumer-grade computer interfaces and laboratory equivalent research tools with no human factors involved. The consumer-grade based testing that utilized visual stimuli demonstrated significant inaccuracies when compared to the tactile based testing.

The fundamental deficiency of commodity-grade computer interfaces for use in high-fidelity human performance research has been a concern for decades among researchers who value accuracy over simplicity. While there are a number of reports that demonstrate that many researchers have developed specialized laboratory equipment for the accurate measurement of human performance, a large and growing cohort of researchers are more frequently using less specialized equipment. From a high-altitude perspective, it appears that there was a time in the scientific study of human performance when researchers actually built and understood their instruments, tested and calibrated them carefully, knew their strengths and limitations, and factored these into the analysis of their results. But with the advent and ubiquity of low-cost commodity-grade human interface devices and displays designed to give the *perception* of smooth operation while being increasingly simple to use and low in cost, the use of these devices as if they were scientific-grade instruments has become alarmingly widespread, while the understanding of how well they work and how accurate and reliable they are has dwindled significantly. This trend was first pointed out with respect to the use of computer mice by Beringer (1992), more recently described by Plant, Hammond, and Whitehouse (2002 & 2003), then by Plant, Hammond, and Turner in 2004, and then again in 2009 (Plant and Turner), the latter after the use of mobile “smart” devices had begun its exponential rise. In their 2009 paper, Plant and Turner update their earlier findings and observe that the trend had not improved. They note that millisecond *precision* is a very different thing from millisecond *accuracy.* Even with newer human interface technologies, timing accuracy has not enjoyed the same priority and improvements over time as cost reduction. They conclude that, “It is important to note that the fact that hardware and software produce answers that ‘look accurate’ does not mean that those answers are valid.”

In 2016 Plant again emphasizes that millisecond timing accuracy errors are prevalent throughout the psychology literature and that this may contribute to what is now recognized as the “replication crisis”. The replication crisis appears to span most of biomedical research, even in cancer and drug development research (Begley and Ellis, 2012; Prinz et al., 2011) in which careful replication exercises demonstrate that as much as 75% to 89% of academic research, published by the best laboratories in top journals, may simply not be reproducible. Many factors contribute to the replication crisis across medical research. Plant suggests that within the field of psychology, this is likely due in part to hardware and software problems that contribute to timing errors and reproducibility problems between different laboratories. Plant further points out that faster hardware has not improved timing accuracy, rather over the years it has apparently gotten worse, and that most researchers simply do not know what their timing accuracy actually is. Further complicating the issue is the fact that web-based studies have become increasingly common and have introduced several new sources of inaccuracy, including server load and caching of scripts. Plant states, “In sum, accuracy has continued to decrease but our confidence in the equipment and the perception of accuracy has risen as computers have become faster and ubiquitous.”

For a scientific measure to be valid, it must be both accurate and precise. The problem with commodity-grade computer interfaces is that they may be precise while not being accurate, or they may lack both precision and accuracy because they typically introduce constant or variable non-zero timing offset biases that cannot simply be overcome by “taking more samples” and relying upon the central limit theorem as suggested by Ulrich and Giray (Ulrich and Giray, 1989). Using this approach with any form of systematic bias, additional samples will only render a result that is more precisely inaccurate. The paradox faced by researchers is that while modern commodity-grade human computer interface devices and networking increasingly gain the veneer of smooth glitch-less operation, precise timing tasks in the background are compromised in increasingly subtle ways that are more difficult to detect, quantify, predict, and eliminate.

Could improvements to the accuracy and precision of reaction time testing increase the reliability of cognitive assessments? In the authors’ opinion, this is a resounding yes. To address this question, consider the example of mild traumatic brain injury (mTBI), which is just one of many neurological disorders that demonstrate an altered reaction time. Multiple reports have noted the importance of reaction time assessment in monitoring mTBI and concussion (Ruesch, 1944; Van Zomeren and Deelman, 1978; Macflynn et al., 1984; Stuss et al., 1989; Ponsford and Kinsella, 1992; Collins and Long, 1996; Zahn and Mirsky, 1999; Warden et al., 2001; Collins et al., 2003; Sarno et al., 2003; Willison and Tombaugh, 2006; Niogi et al., 2008; Gould et al., 2013; Eckner et al., 2015). More recently, investigators have recognized that reaction time variability is a better indicator for cognitive function than reaction time alone, suggesting that it is much more sensitive to neurological disorders such as concussion (Cole et al., 2018).

The evaluation of individuals who have sustained mild traumatic brain injury has been growing in prominence in the public forum, with much of this debate arising from the widespread inadequacy of the methods commonly used to assess cognitive function and the neurological insults that are caused by mTBI. One of the measures that is commonly obtained by most online cognitive assessment tools is simple reaction time, yet very few of these online assessment tools have the capacity to evaluate the metric accurately, much less the capacity to evaluate reaction time variability. Reaction time variability has a normative range of 10-20 msec, which simply cannot be measured by systems that have variable latency ranges of 84 msec (normative reaction time is in the 200 msec range, so introduction of this amount of error is also significant). Additionally, many of these assessments are performed on multiple computers and operating systems for the same subject, which can lead to errors caused by inconsistencies between different systems. As mentioned above, reaction time variability appears to be a more important measure for mTBI assessment than reaction time, and given the extreme sensitivity of this measure to the timing errors introduced by commodity grade computing, the need for improved accuracy is significant.

In summary, the question that we sought to address was whether or not we, as a scientific community, have obtained and published reaction time data over the past several decades that has introduced latencies and errors to our knowledge base. In our opinion, the results of this study support the idea that the slower reaction times that have been reported in the literature are not the result of growing neurological deficits in the general population but rather are the result of a weakening of the technical standards that many of the published studies adhere to. Thus, despite improvements in overall technology, it is probably correct to assert that the accuracy of reaction time testing, for most contemporary methods, has not improved in over 150 years, and researchers should pay close attention to the methods that they utilize.

## Acknowledgements

Research reported in this article was supported in part by the Office of Naval Research.

## Supplemental Figures: Tables

**Table 1.**
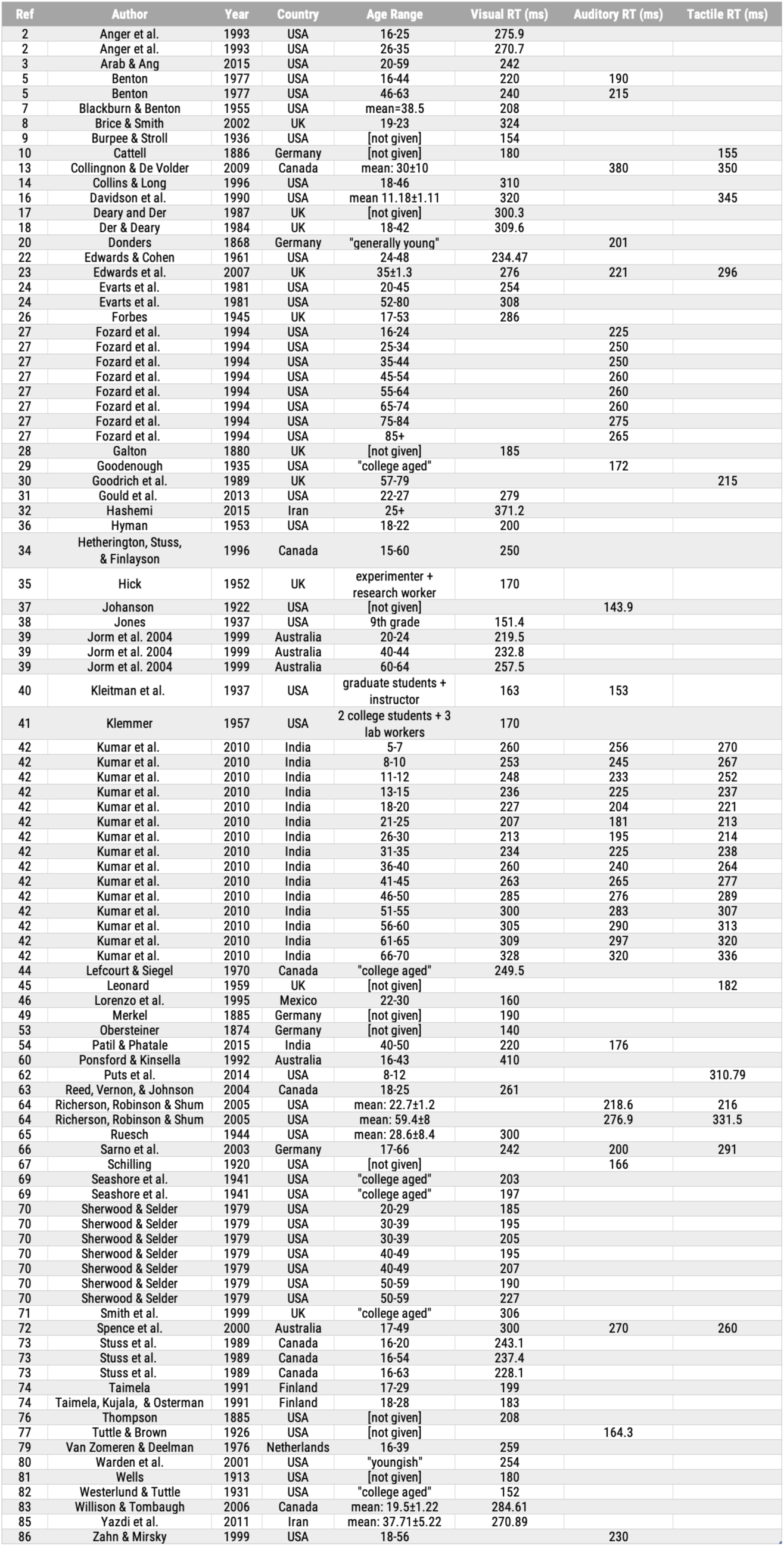
Reaction time values reported in the literature. Values are sorted by year collected, age of subject, and modality of measurement (visual reaction time, auditory reaction time, and tactile reaction time).

| Ref | Author | Year | Country | Age Range | Visual RT (ms) | Auditory RT (ms) | Tactile RT (ms) |
| --- | --- | --- | --- | --- | --- | --- | --- |
| 2 | Anger et al. | 1993 | USA | 16-25 | 275.9 |  |  |
| 2 | Anger et al. | 1993 | USA | 26-35 | 270.7 |  |  |
| 3 | Arab & Ang | 2015 | USA | 20-59 | 242 |  |  |
| 5 | Benton | 1977 | USA | 16-44 | 220 | 190 |  |
| 5 | Benton | 1977 | USA | 46-63 | 240 | 215 |  |
| 7 | Blackburn & Benton | 1955 | USA | mean=38.5 | 208 |  |  |
| 8 | Brice & Smith | 2002 | UK | 19-23 | 324 |  |  |
| 9 | Burpee & Stroll | 1936 | USA | [not given] | 154 |  |  |
| 10 | Cattell | 1886 | Germany | [not given] | 180 |  | 155 |
| 13 | Collingnon & De Volder | 2009 | Canada | mean: 30±10 |  | 380 | 350 |
| 14 | Collins & Long | 1996 | USA | 18-46 | 310 |  |  |
| 16 | Davidson et al. | 1990 | USA | mean 11.18±1.11 | 320 |  | 345 |
| 17 | Deary and Der | 1987 | UK | [not given] | 300.3 |  |  |
| 18 | Der & Deary | 1984 | UK | 18-42 | 309.6 |  |  |
| 20 | Donders | 1868 | Germany | "generally young" |  | 201 |  |
| 22 | Edwards & Cohen | 1961 | USA | 24-48 | 234.47 |  |  |
| 23 | Edwards et al. | 2007 | UK | 35±1.3 | 276 | 221 | 296 |
| 24 | Everts et al. | 1981 | USA | 20-45 | 254 |  |  |
| 24 | Everts et al. | 1981 | USA | 52-80 | 308 |  |  |
| 26 | Forbes | 1945 | UK | 17-53 | 286 |  |  |
| 27 | Fozard et al. | 1994 | USA | 16-24 |  | 225 |  |
| 27 | Fozard et al. | 1994 | USA | 25-34 |  | 250 |  |
| 27 | Fozard et al. | 1994 | USA | 35-44 |  | 250 |  |
| 27 | Fozard et al. | 1994 | USA | 45-54 |  | 260 |  |
| 27 | Fozard et al. | 1994 | USA | 55-64 |  | 260 |  |
| 27 | Fozard et al. | 1994 | USA | 65-74 |  | 260 |  |
| 27 | Fozard et al. | 1994 | USA | 75-84 |  | 275 |  |
| 27 | Fozard et al. | 1994 | USA | 85+ |  | 265 |  |
| 28 | Galton | 1880 | UK | [not given] | 185 |  |  |
| 29 | Goodenough | 1935 | USA | "college aged" |  | 172 |  |
| 30 | Goodrich et al. | 1989 | UK | 57-79 |  |  | 215 |
| 31 | Gould et al. | 2013 | USA | 22-27 | 279 |  |  |
| 32 | Hashemi | 2015 | Iran | 25+ | 371.2 |  |  |
| 36 | Hyman | 1953 | USA | 18-22 | 200 |  |  |
| 34 | Hetherington, Stuss, & Finlayson | 1996 | Canada | 15-60 | 250 |  |  |
| 35 | Hick | 1952 | UK | experimenter + research worker | 170 |  |  |
| 37 | Johanson | 1922 | USA | [not given] |  | 143.9 |  |
| 38 | Jones | 1937 | USA | 9th grade | 151.4 |  |  |
| 39 | Jorm et al. 2004 | 1999 | Australia | 20-24 | 219.5 |  |  |
| 39 | Jorm et al. 2004 | 1999 | Australia | 40-44 | 232.8 |  |  |
| 39 | Jorm et al. 2004 | 1999 | Australia | 60-64 | 257.5 |  |  |
| 40 | Kleitman et al. | 1937 | USA | graduate students + instructor | 163 | 153 |  |
| 41 | Klemmer | 1957 | USA | 2 college students + 3 lab workers | 170 |  |  |
| 42 | Kumar et al. | 2010 | India | 5-7 | 260 | 256 | 270 |
| 42 | Kumar et al. | 2010 | India | 8-10 | 253 | 245 | 267 |
| 42 | Kumar et al. | 2010 | India | 11-12 | 248 | 233 | 252 |
| 42 | Kumar et al. | 2010 | India | 13-15 | 236 | 225 | 237 |
| 42 | Kumar et al. | 2010 | India | 18-20 | 227 | 204 | 221 |
| 42 | Kumar et al. | 2010 | India | 21-25 | 207 | 181 | 213 |
| 42 | Kumar et al. | 2010 | India | 26-30 | 213 | 195 | 214 |
| 42 | Kumar et al. | 2010 | India | 31-35 | 234 | 225 | 238 |
| 42 | Kumar et al. | 2010 | India | 36-40 | 260 | 240 | 264 |
| 42 | Kumar et al. | 2010 | India | 41-45 | 263 | 265 | 277 |
| 42 | Kumar et al. | 2010 | India | 46-50 | 285 | 276 | 289 |
| 42 | Kumar et al. | 2010 | India | 51-55 | 300 | 283 | 307 |
| 42 | Kumar et al. | 2010 | India | 56-60 | 305 | 290 | 313 |
| 42 | Kumar et al. | 2010 | India | 61-65 | 309 | 297 | 320 |
| 42 | Kumar et al. | 2010 | India | 66-70 | 328 | 320 | 336 |
| 44 | Lefcourt & Siegel | 1970 | Canada | "college aged" | 249.5 |  |  |
| 45 | Leonard | 1959 | UK | [not given] |  |  | 182 |
| 46 | Lorenzo et al. | 1995 | Mexico | 22-30 | 160 |  |  |
| 49 | Merkel | 1885 | Germany | [not given] | 190 |  |  |
| 53 | Obersteiner | 1874 | Germany | [not given] | 140 |  |  |
| 54 | Patil & Phatale | 2015 | India | 40-50 | 220 | 176 |  |
| 60 | Ponsford & Kinsella | 1992 | Australia | 16-43 | 410 |  |  |
| 62 | Puts et al. | 2014 | USA | 8-12 |  |  | 310.79 |
| 63 | Reed, Vernon, & Johnson | 2004 | Canada | 18-25 | 261 |  |  |
| 64 | Richerson, Robinson & Shum | 2005 | USA | mean: 22.7±1.2 |  | 218.6 | 216 |
| 64 | Richerson, Robinson & Shum | 2005 | USA | mean: 59.4±8 |  | 276.9 | 331.5 |
| 65 | Ruesch | 1944 | USA | mean: 28.6±8.4 | 300 |  |  |
| 66 | Sarno et al. | 2003 | Germany | 17-66 | 242 | 200 | 291 |
| 67 | Schilling | 1920 | USA | [not given] |  | 166 |  |
| 69 | Seashore et al. | 1941 | USA | "college aged" | 203 |  |  |
| 69 | Seashore et al. | 1941 | USA | "college aged" | 197 |  |  |
| 70 | Sherwood & Selder | 1979 | USA | 20-29 | 185 |  |  |
| 70 | Sherwood & Selder | 1979 | USA | 30-39 | 195 |  |  |
| 70 | Sherwood & Selder | 1979 | USA | 30-39 | 205 |  |  |
| 70 | Sherwood & Selder | 1979 | USA | 40-49 | 195 |  |  |
| 70 | Sherwood & Selder | 1979 | USA | 40-49 | 207 |  |  |
| 70 | Sherwood & Selder | 1979 | USA | 50-59 | 190 |  |  |
| 70 | Sherwood & Selder | 1979 | USA | 50-59 | 227 |  |  |
| 71 | Smith et al. | 1999 | UK | "college aged" | 306 |  |  |
| 72 | Spence et al. | 2000 | Australia | 17-49 | 300 | 270 | 260 |
| 73 | Stuss et al. | 1989 | Canada | 16-20 | 243.1 |  |  |
| 73 | Stuss et al. | 1989 | Canada | 16-54 | 237.4 |  |  |
| 73 | Stuss et al. | 1989 | Canada | 16-63 | 228.1 |  |  |
| 74 | Taimela | 1991 | Finland | 17-29 | 199 |  |  |
| 74 | Taimela, Kujala, & Osterman | 1991 | Finland | 18-28 | 183 |  |  |
| 76 | Thompson | 1885 | USA | [not given] | 208 |  |  |
| 77 | Tuttle & Brown | 1926 | USA | [not given] |  | 164.3 |  |
| 79 | Van Zomeren & Deelman | 1976 | Netherlands | 16-39 | 259 |  |  |
| 80 | Warden et al. | 2001 | USA | "youngish" | 254 |  |  |
| 81 | Wells | 1913 | USA | [not given] | 180 |  |  |
| 82 | Westerlund & Tuttle | 1931 | USA | "college aged" | 152 |  |  |
| 83 | Willison & Tombaugh | 2006 | Canada | mean: 19.5±1.22 | 284.61 |  |  |
| 85 | Yazdi et al. | 2011 | Iran | mean: 37.71±5.22 | 270.89 |  |  |
| 86 | Zahn & Mirsky | 1999 | USA | 18-56 |  | 230 |  |

**Table 2.** Latencies of response to simulated reaction time testing.

| Modality | Average Latency | Range of Latencies |
| --- | --- | --- |
| Visual RT + Touchscreen | 399 ± 16.3 | 348 - 432 |
| Visual RT + Mouse | 80.1 ± 8.0 | 59 - 104 |
| Tactile RT + Mouse | 31.7 ± 2.6 | 27 - 37 |
| Tactile RT + BG | 5.6 ± 0.25 | 4.8 - 6.1 |

